# Robust prediction of drug combination side effects in realistic settings

**DOI:** 10.1101/2025.10.16.682750

**Authors:** Rubén Jiménez, Alberto Paccanaro

**Affiliations:** Escola de Matemática Aplicada, Fundação Getúlio Vargas, Rio de Janeiro, Brazil; Department of Computer Science, Centre for Systems and Synthetic Biology, Royal Holloway University of London, Egham, UK

## Abstract

Side effects caused by drug combinations pose a major challenge in healthcare. Knowledge of these side effects is limited because often they are not detected in clinical trials, which typically involve a restricted number of participants and tested drug combinations. We introduce DCSE (Drug Combinations Side Effects), a novel machine learning method for predicting polypharmacy side effects. DCSE learns latent signatures for drugs, drug pairs, and side effects to predict the probability that a side effect occurs in a given drug combination. We first evaluate its performance in the commonly adopted experimental settings in the literature. Then, a key contribution of this paper is the introduction of more realistic experimental settings that incorporate warm-start and cold-start scenarios under a prospective evaluation. Here, we attempt to predict side effects reported between 2009 and 2014 after training only on data available prior to that period. Our results indicate that DCSE consistently outperforms state-of-the-art methods, demonstrating its robustness and efficacy in real-world applications.

**Author summary:** Understanding the risks of combining medications is a central problem in healthcare. Clinical trials rarely capture the full range of adverse reactions, and many side effects are identified only after drugs reach the market, leaving many possible drug combinations insufficiently tested. Computational approaches can help fill this gap by predicting which combinations are likely to cause specific side effects. Our approach works by learning signatures for both drugs and side effects, which can be thought of as representations that describe how drugs act and what is required to trigger a side effect. A key challenge is how to combine the signatures of two drugs into one representation for the pair. We show that simple linear models fail to capture the complexity of drug interactions, so we use a deep neural network to combine them in a non-linear way. Our tests, which use reports collected at different points in time to mirror how information about drug pairs becomes available, show that our approach is useful for monitoring risks after drugs have been approved and for exploring new uses of pairs of existing drugs.

## Introduction

Many human diseases arise from highly complex biological processes that resist the influence of a single drug. Polypharmacy, the concurrent use of multiple medications, is a common strategy to overcome such limitation [1, 2]. However, a major drawback of polypharmacy therapies is the increased risk of side effects triggered by drug combinations [3,4] that, in the U.S. alone, are affecting over 15% of the population and costing over 170 billion annually for treatment [5]. Healthcare providers often lack awareness of the clinical risks associated with most drug combinations. Incomplete knowledge about these side effects stems from the limited sample size of clinical trial participants, that restricts the ability to detect less common side effects [2], along with the infeasibility of experimentally testing all possible drug combinations during clinical trials.

In the past decade, a wealth of information relating to interactions between pairs of drugs, also known as Drug-Drug interactions (DDI), has become publicly available [6,7]. This allowed the development of several computational methods to train machine learning models for predicting adverse side effects caused by DDIs. Early methods were not specific to individual side effects, instead focusing primarily on predicting the probability that two drugs would interact, causing any side effect [8–11]. More recently, machine learning methods have shown promising results in identifying the specific side effects caused by drug pairs. Among these are graph-based methods, such as Decagon, the pioneering work by Zitnik et al. [5,12,13], that models DDI data as a multi-relational knowledge graph, where each side effect is treated as a distinct edge type. A graph convolutional network is then trained to perform multi-relational link prediction between drugs. Traditional graph-based methods, such as DeepWalk [14], and tree-based models like XGBoost [15], have also been widely adopted as baselines for evaluating polypharmacy side effect prediction methods. A different approach, DeepDDI, presented by Ryu et al. [16], represents each drug with a structural similarity profile and concatenates them to form a drug pair input to a deep feedforward neural network that predicts specific adverse events. Nyamabo et al. [17,18] introduced GMPNN, a message-passing neural network that learns representations of molecular substructures from drug graphs and combines them to predict side effects for drug pairs. They further demonstrate the effectiveness of GMPNN in predicting interactions involving drugs with limited or no prior information (cold-start scenarios).

In this paper, we present DCSE (Drug Combinations Side Effects, pronounced “dixie”), a method for predicting the probability of adverse side effects caused by DDIs. Our approach learns signatures (latent feature vectors) separately for each drug and then combines them nonlinearly using a deep neural network to obtain the signature for the drug pair. Prediction is achieved through the linear combination of drug pairs and side effect signatures, and it can be interpreted as the probability of the side effect being associated with the drug pair. We first show that DCSE outperforms state-of-the-art methods in the standard experimental settings adopted in the literature. We then conduct experiments in a more realistic setting, aiming to predict the probability of associations that were unknown at the time the training set was created. In our prospective experiments, we distinguish two scenarios: warm-start, where we predict side effects for drug pairs for which some side effects are already known; and cold-start, where we predict side effects for drug pairs for which no side effects are known. Our results demonstrate that DCSE consistently delivers robust performance across both standard and more realistic scenarios, outperforming competing methods.

## Results

### The DCSE model

We recently showed [19] that drugs and side effects can be represented as vectors (signatures) in a low-dimensional space, such that the frequency class ^1^ of the side effect for a single drug can be estimated using the dot product of their corresponding vectors. Figure 1a shows a representation of this idea for the drug “metformin” and the side effect “diarrhea”. The signatures can be interpreted as representing the states of components in a biological system: the elements of a drug signature reflect the intensity of the drug’s action on these components, while the elements of a side effect signature indicate which components need to be affected to trigger a specific side effect.

**Fig 1.**
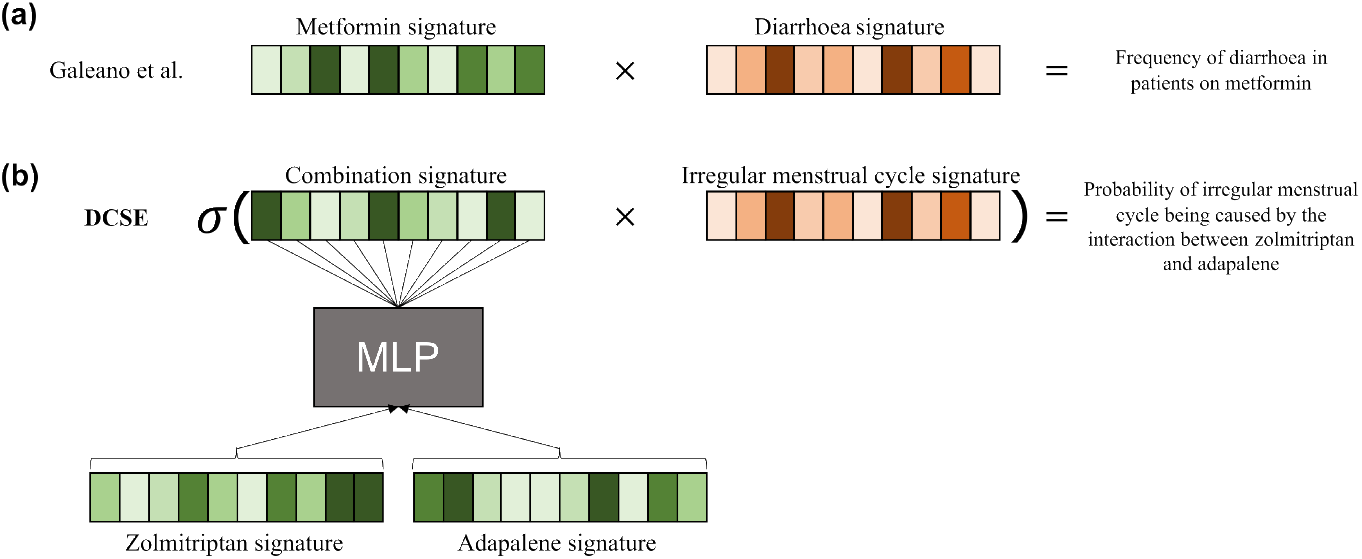
The approaches by Galeano et al. (2020) and DCSE. **(a)** Galeano et al. [19] showed that the frequency of diarrhoea in patients on metformin can be obtained by the dot product of the signature for metformin and the signature for diarrhoea. **(b)** In DCSE, the learned signatures for zolmitriptan and adapalene are combined using an MLP to obtain the combination signature. The probability of an irregular menstrual cycle being caused by this pair is obtained by the logistic function of the product of the drug combination signature and the side effect signature.

The idea of representing the state of a system using signatures could potentially be applied to the problem of predicting the probability of side effects caused by drug pairs, provided we could learn signatures representing the intensity of the action of a drug pair on the system components. Can we combine the signatures of two drugs to derive a signature for the pair? Initially, we explored learning a linear combination of the signatures of individual drugs. As expected, our experiments demonstrated that a linear model cannot effectively capture the complexity of combining drug signatures (see *S1 Text*). This result aligns with the well-established understanding that the effects of drug combinations are highly non-linear [21].

Consequently, we opted to use a deep neural network to nonlinearly combine the signatures of the drugs. Figure 1b illustrates this approach, that we call DCSE: the learned drug signatures for zolmitriptan and adapalene are combined through a Multi-Layer Perceptron (MLP) to generate the signature for the pair. The probability of the side effect “irregular menstrual cycle” being caused by the zolmitriptan-adapalene pair is then calculated as the logistic function of the dot product between their corresponding signatures.

An overview of the complete DCSE architecture is shown in Figure 2, and a formal description of the system can be found in *Methods*. The input comprises three one-hot encoding vectors, representing the identity of the two drugs, drug *i* and drug *j*, and the side effect *l*. Above the input layers are the embedding layers, fully connected layers that linearly project the one-hot encodings to dense vectors, which are the drug (**w**_**i**_ and **w**_**j**_) and side effect (**h**_**l**_) signatures. To ensure consistent representations, the weights in the layer *W* are shared between the two drug inputs, so that a drug has the same signature, regardless of its position in the pair. The signatures for the two drugs are concatenated and passed through a fully-connected MLP architecture, which we refer to as the combination layers. The output of the combination layers, **c**_*i,j*_, is the signature of the combination of drugs *i* and *j*.

**Fig 2.**
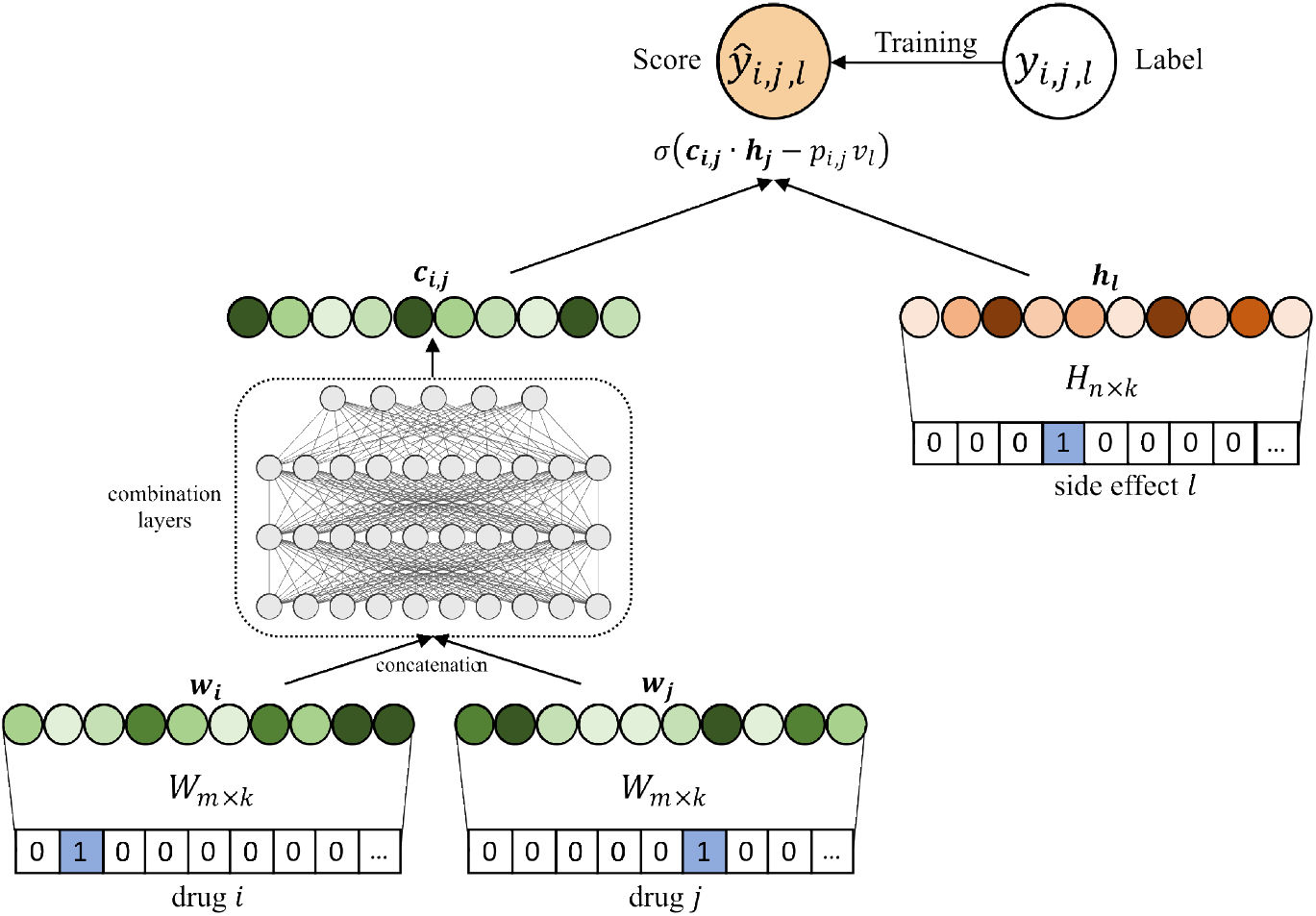
Architecture of DCSE. The model learns signatures for drugs and side effects, *w*_*i*_, *w*_*j*_, and *h*_*l*_, respectively, using an embedding layer. The drug signatures are then combined through an MLP to obtain the combination signature, *c*_*i,j*_. The final prediction *ŷ*_*i,j,l*_ is obtained by the dot product between the signatures of the combination and the side effect and then subtracted by the product of the drug combination and side effect biases, *p*_*i,j*_ and *v*_*l*_, respectively.

When learning from association data, it is essential to account for inherent biases, such as prescription bias — where some drugs are prescribed more frequently — and the higher prevalence of certain side effects in the population [21,22]. DCSE addresses this by explicitly modeling biases for individual drugs, combinations, and side effects. Our approach learns a vector of drug biases, where each element is the bias of a drug, and a vector of side effect biases, where each element is the bias of a side effect. Biases for drug pairs (*p*_*i,j*_) are learned by nonlinearly combining the biases of the individual drugs using a small MLP (not shown in the figure). Our experiments demonstrate that these biases capture systematic tendencies in the data, such as the “popularity” of specific drug pairs and side effects (see S1 Text).

The predicted probability of side effect *l* for the combination of drugs *i* and *j* is calculated by first computing the dot product between the signature of the drug pair and the signature of the side effect. To this value, a bias term is then subtracted (see *Methods)*, that is calculated by multiplying the bias for the drug pair *p*_*i,j*_ and the bias of the side effect *v*_*l*_. This is then transformed using a sigmoid function, which maps it to the probability of the side effect being associated with the drug pair.

All the weights in this model are learned using the Twosides [6] database of polypharmacy side effects, which provides statistically significant associations between side effects and drug pairs (see *Methods*). All parameters are learned simultaneously, including the single drug and side effect signatures, the weights of the deep neural network, and the biases.

### Evaluation

We first evaluate DCSE using widely adopted experimental settings in the DDI prediction literature. Next, we present a discussion on the limitations inherent in these settings and propose more realistic evaluation procedures. Our novel evaluations are carried out prospectively: we attempt to predict side effects reported between 2009 and 2014 after training only on data available prior to this period. Within this framework, we also distinguish between scenarios where drug pairs in testing are encountered during training (referred to as warm-start) and scenarios where testing involves drug pairs that are entirely new (referred to as cold-start).

#### Conventional experimental settings

We selected two evaluation procedures commonly adopted in the literature: the first proposed by Zitnik et al. [5] and the second by Nyamabo et al. [17]. The Zitnik et al. procedure begins by preprocessing the Twosides dataset to ensure a minimum amount of information for each side effect. Specifically, only side effects occurring in at least 500 drug combinations are retained. For each side effect, associations are randomly divided into training (80%), validation (10%), and testing (10%) sets. Negative samples are selected for training by substituting one of the drugs in a positive association. In the training set, the replacement drug is selected according to a sampling distribution defined empirically by Mikolov et al. [23]. In the testing set, the drugs are sampled randomly from a uniform distribution. Both training and testing phases are conducted under a balanced setting, i.e., with a ratio of positives to negatives of 1: 1.

The Nyamabo et al. setup is based on the procedure by Zitnik et al., but with a few variations: the percentages for the splits are adjusted to 60% for training, 20% for validation, and 20% for testing; negative samples are selected using a different sampling distribution, defined by Wang et al. [24], for both training and testing; and the entire process is repeated three times, resulting in three stratified randomized folds. Again, both training and testing phases are conducted under a balanced setting.

We compared the performance of DCSE against DeepWalk [14], XGBoost [15], DeepDDI [16], Decagon [5], and GMPNN [17] (these algorithms are described in the *Methods* section). These baseline methods and state-of-the-art competitors were selected because they (1) are the representative methods of the most commonly adopted experimental setups in the literature; (2) encompass a wide range of machine learning methods; (3) integrate various sources of information as input features for the drugs; and/or (4) work for both warm and cold-start scenarios. DeepWalk and XGBoost do not support multi-label classification and required training one model independently for each side effect, as already implemented by Zitnik et al. [5]. In the case of DeepDDI, since in this setting different drug pairs have different numbers of side effects, we also trained one model independently for each side effect.

The results of these evaluations are presented in Table 1. We can observe that DCSE’s performance is consistent across the two evaluation procedures. Our results show that the signatures learned by DCSE are descriptive and effective, outperforming methods with more complex architectures, such as Decagon, which relies on graph convolutional networks, and GMPNN, which uses gated message passing neural networks. It is also important to note that the superior performance of end-to-end methods (DCSE, Decagon, and GMPNN) compared to the feature-based approaches (DeepWalk, XGBoost, and DeepdDI). This suggests that for the complex task of predicting side effects for drug combinations, methods greatly benefit from learning directly from the association data. The reduced performance of DeepWalk, XGBoost, and DeepDDI could also be due to the fact that training one model per side effect prevents information from being shared across side effects. As a result, these methods cannot fully capture the interdependence between drugs and side effects.

**Table 1.** Comparative benchmark in conventional experimental evaluation. For the Nyamabo et al. setting, the mean and standard deviation across the three folds are shown. The best performance is shown in bold. † Results taken from Zitnik et al. [5].

| Method | *Zitnik et al. Evaluation* |  | Method | *Nyamabo et al. Evaluation* |  |
| --- | --- | --- | --- | --- | --- |
|  | AUROC | AUPRC |  | AUROC | AUPRC |
| DeepWalk | 71.51 | 70.9 | DeepWalk | 70.13 ± 0.1 | 69.02 ± 0.11 |
| XGBoost | 80.12 | 81.15 | XGBoost | 78.42 ± 0.13 | 78.84 ± 0.14 |
| DeepDDI | 62.37 | 61.04 | DeepDDI | 60.44 ± 0.07 | 59.81 ± 0.06 |
| Decagon† | 87.23 | 83.2 | GMPNN | 90.07 ± 0.12 | 87.24 ± 0.12 |
| **DCSE** | **91.71** | **87.24** | **DCSE** | **92.44 ± 0.03** | **89.67 ± 0.02** |

#### Our proposed prospective experimental settings

A concern with evaluating system performance in the above experimental settings is the reliance on a balanced test set, which does not accurately reflect the true class distribution. For instance, in the Twosides dataset, among the 83,657,414 possible associations between 63,473 drug pairs with at least one side effect association and 1,318 side effects, only 4,651,131 are known associations (positives), while the remaining 79,006,283 are unknown (negatives). Therefore, negatives account for 94.44% of all possible associations, a distribution that is far from balanced. We also notice that not all negative associations are equally difficult to predict. One can argue that drug pairs for which no side effects are currently known are more likely to be acting on completely different biological systems and are therefore very unlikely to cause any side effects.

Thus, a negative association between drug pairs with no known associations should be generally easier to predict. For the 645 drugs in Twosides, taking all possible combinations two at a time results in 207,690 possible drug pairs. Of these, 144,217 have no known side effects at all, and this amounts to a total of 190,078,006 negative associations that are easier to predict. In contrast, for the 79,006,283 associations from the 63,473 drug pairs that have some known associations, predicting the absence of a specific side effect is generally more difficult. In conventional experimental settings, the procedure for sampling negative examples does not control for this distinction, making it highly likely that easier pairs dominate the negative samples.

To address these issues, we introduce a different experimental setting for evaluating performance on this problem. First, we do not balance the testing datasets. Second, we focus only on the more challenging negative associations by taking the 79,006,283 unknown observations between the drug pairs and side effects in Twosides.

We perform our experiments in a prospective setting, where the goal is to predict new side effects reported in an updated version^2^ of Twosides, which includes reports from 2009 to 2014, while training on the original version with reports up to 2009. We refer to these datasets as Twosides-2014 and Twosides-2009, respectively. These tests provide a more realistic assessment of the method’s performance in real-life scenarios, as they account for the chronological order in which side effects are discovered. This aspect is important because it has been noted that associations between side effects are often related [25].

We also distinguish between two distinct scenarios: one in which some side effects are already available for the drug combinations (warm-start), and another in which no prior data exists (cold-start). These two scenarios reflect different real-world situations. Warm-start experiments are relevant for post-marketing pharmacovigilance, which involves the identification, evaluation, and mitigation of drug-related issues after a drug or drug combination has been released to the market [26]. Cold-start experiments are relevant for drug repurposing, where identifying potential side effects can help accelerate the approval process for new drug combinations [1,27].

Finally, we assess DCSE’s performance by benchmarking it against DeepWalk, XGBoost, and DeepDDI. For DeepWalk and XGBoost, we again follow the procedure described by Zitnik et al. [5], and train one model per side effect. For DeepDDI, in this setting, we can frame the problem as a multi-label classification problem and follow the original procedure described by Ryu et al. [16]. Decagon and GMPNN were excluded from this evaluation due to technical constraints in the source code (see *Methods*). Our focus here is not on an exhaustive comparison, but on evaluating performance in more realistic prospective scenarios.

#### Warm-start evaluation

Our warm-start evaluation is shown in Figure 3. We train a model using Twosides-2009 (the middle matrix in the figure) and use Twosides-2014 (the matrix on the left) as the test set. Twosides-2014 contains 3,642,690 new associations not present in the training data. All entries labeled as negatives in Twosides-2009 are used in testing, and evaluation is performed by ranking these entries and identifying which of them were confirmed as positives in Twosides-2014 (green cells). The gray cells correspond to the positive associations during training and, hence, are not considered during testing. This evaluation protocol conceptually resembles matrix completion, where predictions are made over a fixed set of unknown entries in a partially observed matrix, in contrast to conventional settings where both positives and negatives differ between training and testing.

**Fig 3.**
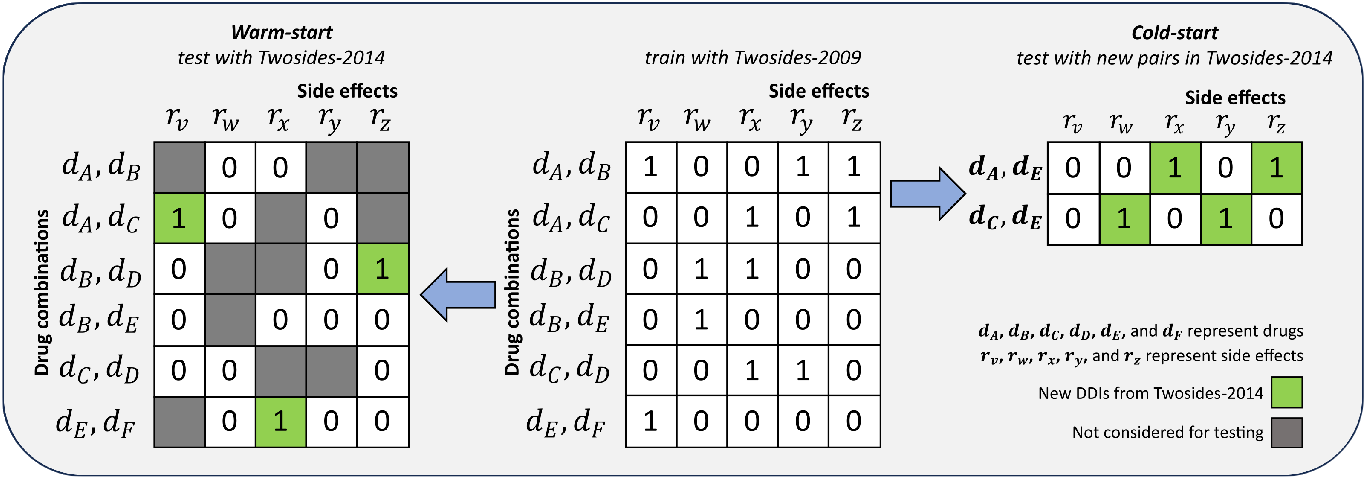
Prospective Evaluation Scenarios. The center of the figure shows our training dataset, Twosides-2009, with drug combinations on the rows and side effects on the columns. This entire dataset is used to train the model for both warm-start and cold-start scenarios. In warm-start (left-side), the model is tested on new associations from Twosides-2014 involving the same drug combinations from Twosides-2009. Cold-start (right-side), testing is done on new drug combinations introduced in Twosides-2014. Note that while pairs *d*_*A*_, *d*_*E*_ and *d*_*C*_, *d*_*E*_ were not present during training, the single drugs *d*_*A*_, *d*_*E*_, and *d*_*C*_ were present but in combination with other drugs.

The results of this evaluation are presented in the “Warm-start” columns in Table 2. The shift to an unbalanced testing set, with *∼*1: 20 positive-to-negative ratio, is reflected in the sharp drop in AUPRC across all methods, as this metric is highly sensitive to class imbalance [28]. DCSE’s performance indicates that the signatures learned for drug combinations effectively capture their interplay with the different side effects and can generalize well to predict new associations. For DeepDDI, we observe a notable improvement in performance compared to the setting where separate models were trained for each side effect. This gain can be attributed to the unified multi-label training setup, which allows the model to exploit dependencies between side effects.

**Table 2.** Performance results in our prospective evaluations. Best performance is shown in bold.

|  | Prospective |  |  |  |
| --- | --- | --- | --- | --- |
|  | Warm-start |  | Cold-start |  |
|  | AUROC | AUPRC | AUROC | AUPRC |
| DeepWalk | 79.70 | 19.62 | 63.42 | 4.69 |
| XGBoost | 69.56 | 15.27 | 56.89 | 4.34 |
| DeepDDI | 84.71 | 18.34 | 85.35 | 13.23 |
| DCSE | **85.87** | **21.85** | **88.71** | **15.34** |

#### Cold-start evaluation

The cold-start setting amounts to a *de-novo* prediction of side effects for new drug combinations. The testing set corresponds to the new drug pairs in Twosides-2014 that are not present in Twosides 2009, while ensuring that the drugs forming these pairs appeared during training. The composition of the test set can be observed in the **“Cold-start”** matrix of Fig 3, where the pair composed of drugs *d*_*A*_ and *d*_*E*_ is absent from the training data, but both drugs are present individually in pairs involving other drugs in the training set (Twosides-2009). For our negative associations, we use all unknown drug pair-side-effect observations corresponding to these new pairs.

The results of this evaluation are presented in the “Cold-start” columns in Table 2. In this case, we have a *∼*1: 33 positive-to-negative ratio. It is important to note that DCSE can make these predictions because it learns signatures for individual drugs. As shown in Figure 2, each drug is passed through a shared embedding module that produces its signature. The individual drug signatures are then passed through the combination layers to generate a signature for the drug combination. This allows DCSE to produce signatures even for pairs not seen during training, as long as both drug signatures are available. Among the competing methods, DeepDDI and XGBoost rely on external features as input to their models. Our results show that, even in the absence of external information about single drugs, the signatures learned by DCSE are highly informative of the underlying biological system. Additionally, it suggests that the MLP can generalize to generate informative combination signatures for previously unseen pairs.

## Discussion

We presented DCSE, a novel method for the prediction of polypharmacy side effects. DCSE learns signatures for drugs, combinations, and side effects to infer an association between a pair of drugs and a given side effect. The work by Galeano et al. showed that drugs and side effects can be represented as vectors in a low-dimensional space (signatures), such that the *frequency* of a side effect for a given drug is given by the dot product between their corresponding signatures. This work extends a similar reasoning to the case of drug combinations. We show that the *probability* of a side effect for a given drug combination can be approximated by the logistic of the dot product between their signatures. We also showed that combining individual drug signatures is a highly nonlinear problem, and we developed an approach to learn signatures for pairs of drugs. We examined the role of the bias terms in DCSE and found a strong correlation with the popularity of drug pairs and side effects, as measured by their frequency in the training data. This suggests that the biases capture general prevalence patterns, allowing the model to learn more biologically meaningful signatures (see *S1 Text* for details). DCSE learns signatures for drug combinations solely from interaction data; thus, our work shows that state-of-the-art performance can be achieved by only using information about known interactions between pairs of drugs. We believe that these results can be further improved by integrating external information, such as chemical and substructural information about drugs.

We have shown the significant performance improvement of DCSE over representative state-of-the-art methods in the most widely adopted experimental settings in the literature. We have shown the limitations of these settings and introduced more realistic evaluations, where we used data from Twosides-2009 to make predictions of polypharmacy side effects introduced in Twosides-2014. Our testing sets ensure that the distribution of the classes resembles the true data distribution observed in real-life scenarios more closely. We contend that these experiments offer a more authentic evaluation by closely resembling how side effects are identified in the patient population. We also distinguished between warm and cold-start scenarios, where drug pairs in testing are either encountered during training or are completely new. DCSE consistently shows higher performance across the different scenarios, indicating that the signatures for the drugs and the combinations can effectively capture the interplay with the different side effects and generalize well to new associations.

Galeano et al. showed that their learned signatures offer interpretability in terms of biological information about drug activity at the anatomical and molecular level [19]. We attempted to derive a similar biological interpretation for the signatures learned by DCSE. Our first step was to ensure the stability of these signatures across different model runs [29,30]. However, we encountered challenges with the reproducibility of the signatures learned by DCSE. Nevertheless, DCSE’s predictive performance remains consistent across various random initializations, as indicated by the low standard deviations shown in Table 1.

It is important to acknowledge certain limitations of our approach. Firstly, DCSE is unable to generate *de-novo* predictions for drug combinations formed by drugs that did not appear at all during training. Secondly, DCSE is effectively a collaborative filtering approach, and therefore, it relies on the availability of very diverse association data to learn meaningful signatures. For this reason, it is unlikely to work on datasets such as DrugBank that count only 86 adverse events, that are very few compared to the 1,318 side effects in Twosides. We believe that both these limitations can be addressed by integrating external information, such as chemical and substructural information about drugs, and will tend to become less relevant with the ever-increasing size of available datasets.

## Materials and methods

### Polypharmacy side effects data

Our experiments were carried out using polypharmacy side effects association data from the Twosides database [6], which contains information about 63, 473 drug combinations and 1, 318 side effects. The total number of individual drugs that make up the combinations is 645. In total, there are 4, 543, 686 associations between side effects and drug pairs. Twosides is the result of a data-driven approach that identifies significant drug-event associations from the Adverse Event Reporting System, which collects reports from doctors, patients, and drug companies.

We used two versions of this dataset: Twosides-2009, which includes reports up to and including 2009; and Twosides-2014, which contains reports up to and including 2014. To facilitate further research, we provide our Twosides-2014 datasets for prospective evaluations in 10.6084/m9.figshare.30355195. These datasets were pre-processed to include the new associations for the same drugs and side effects present in Twosides-2009.

### Formal definition of DCSE

Given a dataset containing *m* drugs and *n* side effects, let the input to the model be represented by three vectors, **d**_*i*_, **d**_*j*_, and **r**_*l*_, with a one-hot encoding that determines the identity of the two drugs and the side effect, respectively (see Figure 2). Next is the embedding layer, a fully connected layer that projects the sparse encoding to a dense vector. To ensure consistent representations for all drugs, we use the same layer *W ∈* ℝ ^*m×k*^ for both drug inputs, where *k* is the signature size. For the side effect input, the embedding layer is denoted as *H ∈*ℝ^*n×k*^. Thus, the signatures for drugs and side effects are obtained as follows:

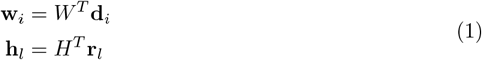

where **d** ∈ℝ^*m×*1^, **r** ∈ℝ^*n×*1^ and **w, h** ∈ℝ^*k×*1^.

The output of the combination layers represents the combination signature and is obtained as follows:

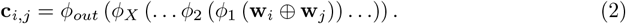

Here ⊕ represents concatenation, *ϕ*_*out*_ and *ϕ*_*x*_ respectively denote the mapping function for the output layer and the *x*_*th*_ hidden layer, where there are *X* hidden layers in total, and **c**_*i,j*_ ∈ ℝ ^*k×*1^ denotes the combination signature of drugs *i* and *j*. It’s important to note that following Galeano et al.’s work [19], *w, h*, and *c* are kept non-negative to favor interpretability of the signatures.

Similarly to how DCSE learns the combination signature, we learn a bias for the combination of drugs by nonlinearly combining the individual biases of the drugs with a small MLP. Specifically, DCSE learns a vector **u** ∈ℝ ^*m×*1^ to represent the drug biases and a vector **v** ∈ ℝ ^*n×*1^ to represent the side effect biases. The biases *u*_*i*_, *u*_*j*_ ∈ ℝ for drugs *i* and *j*, respectively, the side effect bias *v*_*l*_ ∈ ℝ, and the combination bias *p*_*i,j*_ are obtained as follows:

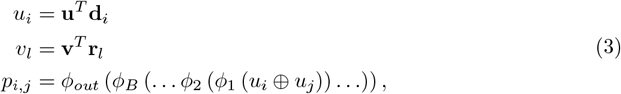

where *B* hidden layers are employed in total for the bias combination MLP. Finally, the predicted score *ŷ*_*i,j,l*_ is obtained by

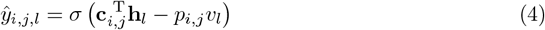

The term 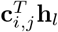 represents the dot product between the combination and side effect signatures, while *p*_*i,j*_*v*_*l*_ denotes the bias term. Here, *σ* represents the sigmoid activation function and maps the output value *ŷ*_*i,j,l*_ to a probability between 0 and 1. Note that since both *ci, j* and *h*_*l*_ are non-negative, their product is also non-negative. To capture the full range of the sigmoid function, the bias term is subtracted from this dot product.

### Implementation Details

All experiments were conducted using an NVIDIA QUADRO RTX 5000 GPU. The model was implemented in PyTorch, and the hyperparameters were tuned using a validation set, following the same procedure described in Section Our proposed prospective experimental settings. The optimal architecture for the combination MLP was a 3-layer neural network, with each hidden layer containing 1, 024 neurons, and a signature size for the drugs, combinations, and side effects of 512. For the bias MLP, the architecture is composed of two hidden layers of 4 neurons each. To ensure the model’s invariance regarding the order of drugs, each input drug is presented in both positions as distinct samples.

The code for the implementation is available at: https://github.com/paccanarolab/DCSE.

### Details about baselines and competitors

- DeepWalk [14]: similar to DCSE, this method learns *k*-dimensional neural features for each drug based on a biased random walk procedure exploring network neighborhoods of nodes. To implement it for our experiments, drug combinations are represented by concatenating learned drug feature representations and used as input to a logistic regression classifier. For each side effect we train a separate classifier. DeepWalk was used as a baseline method in [5].
- XGBoost [15]: to implement this method for our experiments, we follow the procedure detailed by Zitnik et al. [5]. We first construct a feature vector for each drug based on the PCA representation of the drug-target protein interaction matrix and the PCA representation of the side effect profile of each individual drug. The drug pairs are then represented by the concatenation of the corresponding drug feature vectors and used as input to the XGBoost classifier that then attempts to predict the exact side effect for a given drug pair. Similarly to DeepWalk, a separate model is trained for each side effect.
- DeepDDI [16]: in this approach, drugs are represented by a structural similarity profile that contains pairwise structural similarity scores obtained from the comparison between the input drug and all the FDA-approved drugs. The features for each drug are concatenated to represent the drug pair and then fed to a deep neural network to predict the side effect associations. In the experiments following the conventional settings (Section *Conventional experimental settings*), DeepDDI was trained separately for each type of side effect with a single neuron as output, indicating the probability of the side effect occurring for the input drug pair. Later in our proposed experimental settings (Section Our proposed prospective experimental settings), DeepDDI was adjusted – and framed as a multi-label classification problem – to output a probability for all possible side effects given one drug pair, i.e., having the same number of output neurons as side effect types in the dataset.
- Decagon [5]: this approach first constructs a multimodal graph of protein-protein interactions, drug-protein target interactions, and polypharmacy side effects, which are represented as drug-drug interactions, where each edge has multiple types corresponding to all the different side effects. Decagon consists of a graph convolutional neural network for multi-relational link prediction in multimodal networks. As we were unable to run the provided code, we compared our method to Decagon by replicating the same experimental conditions (Section Conventional experimental settings). We were unable to run Decagon’s code base, and therefore, it was excluded from the different evaluation settings as it would require substantial modifications to the original implementation. To still compare fairly against this highly representative method, we include a comparison of DCSE against them using their proposed evaluation methodology.
- GMPNN [17]: uses a Gated Message Passing Neural Network (GMPNN), which learns chemical substructures with different sizes and shapes from the molecular graph representations of drugs for DDI prediction for a pair of drugs. The final prediction is based on the interactions between pairs of their learned substructures. We were able to run the provided code for this competitor, however, only for their experimental setting. Given the challenges in adjusting the code for different experimental settings, we limited the comparison of DCSE to the Nyamabo et al. evaluation. (Section Conventional experimental settings).

## Supporting information

**S1 Text. Supplementary material**. This file includes additional experiments, definitions, and figures.

## Footnotes

1 As defined by the World Health Organisation (WHO) - Council for International Organizations of Medical Sciences (CIOMS) [20]

2 https://nsides.io/

